# The Ca^2+^-binding protein CSE links Ca^2+^-signaling with cell-cell communication in multicellular cyanobacteria

**DOI:** 10.1101/2025.03.26.645587

**Authors:** Teresa Müller, Felicia M. Kleusberg, Karolina Roganowicz, Gregor L. Weiss, Murray Coles, Khaled A. Selim

**Affiliations:** Interfaculty Institute of Microbiology and Infection Medicine, Cluster of Excellence ’Controlling Microbes to Fight Infections’, Tübingen University, 72076 Tübingen, Germany; Department of Protein Evolution, Max Planck Institute for Biology, 72076 Tübingen, Germany; Institute of Molecular Biology and Biophysics, Department of Biology, ETH Zürich, 8093 Zürich, Switzerland; Microbial Biochemistry Group, Institute of Phototrophic Microbiology, Heinrich-Heine University Düsseldorf, Universitätsstraße 1, 40225 Düsseldorf, Germany

**Keywords:** Ca^2+^-signaling, Bacterial septal junctions, Nanopore arrays, Ca^2+^-sensor protein, Cyanobacterial multicellularity, Cell-cell communication machinery

## Abstract

Because of their multicellular lifestyle, filamentous cyanobacteria have evolved sophisticated cell-cell communication machinery to exchange, synchronize, and coordinate the efforts of individual cells in the filament. Analogous to gap junctions that were regarded as a purely Eukaryotic feature, multicellular cyanobacteria were found also to coordinate their cell-cell communication via septal junctions (SJs). However, the molecular signals that regulate the cell-cell communication machinery and SJs assembly are largely unknown. Lately, Ca^2+^- signaling has been implicated in regulating cell junctions in neurons. We recently discovered a new Ca^2+^-sensor protein, CSE, exclusively found in multicellular cyanobacteria. Here, we investigated CSE as a potential link between intracellular Ca^2+^-signaling and cell-cell communication. We solved the solution NMR structure of CSE in its Ca^2+^-bound state and revealed that CSE acts as Ca^2+^-buffer protein. Using cryo-electron tomography, we showed that CSE is not only essential for Ca^2+^ homeostasis, but also mediates cell-cell communication via regulating the formation of nanopores — a necessary precursor of SJs — as Δ*cse* mutant shows a strong reduction in the number of both nanopores and SJs. This determines for the first time Ca^2+^-signaling as a novel mechanism controlling cell-cell communication and establishes CSE as a key player regulating cyanobacterial multicellularity. This highlights also Ca^2+^-signaling as a common conserved principle for regulating cell junctions between organisms that deviated billion years ago.

## Introduction

Calcium (Ca^2+^) ions have the distinction of being both essential and toxic to living cells. Their toxicity stems from the ability to bind and precipitate intracellular phosphate ions, as well as on their negative effect on ribosome functions (Akanuma et al., 2014; Clapham, 2007; Weiss et al., 1973). Therefore, the absolute cellular Ca^2+^ concentrations must be held at nanomolar levels. To achieve this, free calcium ions are either excreted via channels or actively transported through pumps. Further, specific proteins bind calcium ions to buffer the free intracellular Ca^2+^ concentration. This tight control in turn leads to their essential role as a second messenger in a myriad of regulatory processes. Details of these systems have been elucidated over many years. This is particularly true for eukaryotes (Lehne et al., 2022), but some prokaryotic systems are now receiving more attention (Agaras et al., 2023; Burch-Konda et al., 2024; King et al., 2020; Kolodkin-Gal et al., 2023). An example is cyanobacteria, which share common ancestors with plant chloroplasts and have thus been used as models for studying photosynthesis. In plants, vital roles for Ca^2+^-signaling in regulating plant immunity and stress response have been recognized (Aratani et al., 2023; Hagihara et al., 2022; Toyota et al., 2018; Yan et al., 2024). In cyanobacteria, Ca^2+^ signaling is crucial for stress responses to reactive oxygen species (ROS) (Singh et al., 2016), changes in pH (Giraldez-Ruiz et al., 1997), temperature, salt, or osmotic stress (Mantovani et al., 2023; Torrecilla et al., 2000, 2001, 2004). Calcium ions are also implicated in controlling photosynthesis, regulating specific sugar and fatty acid metabolic pathways (Singh et al., 2016, 2017), and fine-tuning carbon/nitrogen metabolism by yet unknown mechanisms (Walter et al., 2016, 2019; Mantovani et al., 2023).

Filamentous cyanobacteria are emerging as model organisms for studying multicellularity, due to their ability to undergo cell differentiation to form heterocysts under nitrogen-limiting conditions (Flores & Herrero, 2010; Maldener & Muro-Pastor, 2010; Muro-Pastor & Maldener, 2019). Heterocysts are specialized cells in filamentous cyanobacteria that fix atmospheric nitrogen to ammonia using nitrogenases, the only family of enzymes known to catalyze this reaction. It has long been known that calcium signaling is essential for proper heterocyst formation (Smith et al., 1987). A transient increase in Ca^2+^ concentration has been observed in pro-heterocysts shortly after nitrogen depletion (Torrecilla et al., 2004; Zhao et al., 2005). The role of Ca^2+^ binding proteins in this process has become later more evident (Mantovani et al., 2023; Walter et al., 2020; Zhao et al., 2005). In the filamentous model organism *Nostoc* sp. PCC 7120 (hereafter *Nostoc*), two calcium binding proteins have been implicated in heterocyst formation: CcbP (<u>C</u>yanobacterial <u>c</u>alcium <u>b</u>inding <u>P</u>rotein; Zhao et al., 2005) and CSE (<u>C</u>a^2+^- <u>S</u>ensor <u>E</u>F-hand; Walter et al., 2019). CcbP is negatively regulated via the heterocyst transcription regulators NtcA and HetR (Forchhammer & Selim, 2020) and appears to be absent in heterocysts (Shi et al., 2006; Zhao et al., 2005). We recently identified CSE as a highly conserved protein in filamentous, nitrogen-fixing cyanobacteria and proved its importance for filament integrity, heterocyst differentiation and photosynthesis (Walter et al., 2019, 2020). The overexpression of *cse* (*cse* OE) impaired both growth and photosynthesis (Walter et al., 2019). In contrast, *cse* deletion (Δ*cse*; encoded by *asr1131*) resulted in strong fragmentation of filaments and negatively influenced heterocyst functions (Walter et al., 2020). Thus, both CSE and CcbP are involved in heterocyst differentiation, highlighting a conserved role for Ca^2+^ signaling in controlling cell differentiation, however, their specific functions and regulatory targets remain unknown.

The role of calcium binding proteins in heterocysts formation hints at a wider role in cell-cell communication (ccc). In multicellular filamentous cyanobacteria, like *Nostoc*, the exchange of intracellular molecules along the filament is crucial both for heterocyst development and diazotrophic growth, as both heterocyst and vegetative cells rely on nutrients from each other (Flores and Herrero, 2010; Maldener et al., 2014). To exchange molecules between sister cells, these metabolites must cross two cytoplasmic membranes and a septal peptidoglycan (PG) disc within the septum between neighboring cells in the filament. Septal junctions (SJs) traverse this PG layer via nanopore arrays drilled by AmiC amidases (N-acetylmurayl-L-alanin amidases; Bornikoel et al., 2017; Lehner et al., 2013; Nürnberg et al., 2015), establishing a connection between the cytoplasm of neighboring cells (Flores et al., 2016; Hoiczyk & Baumeister, 1995; Lehner et al., 2013). SJs are multi-protein complexes, consisting of a septum-spanning tube with a membrane-embedded plug (likely formed by FraD/SepN proteins) at both ends, and a cap covering the plug on the cytoplasmic side (Kieninger et al., 2022; Weiss et al., 2019). The purified PG discs of *Nostoc* can harbor 80-150 nanopores per array. Mutants with disruption in one of the AmiC genes or FraD have a highly reduced number of nanopores and are impaired in cell-cell communication (Bornikoel et al., 2017). Many of those mutants suffer from strong fragmentation phenotypes and impaired heterocyst development, in analogy to Δ*cse* phenotype (Kieninger, 2021; Kieninger & Maldener, 2021; Merino-Puerto et al., 2010; Merino-Puerto et al., 2011; Walter et al., 2020). Thus, there appears to be a strong connection between heterocyst development, filament integrity, and cell-cell communication. However, the molecular signals that control the cell-cell communication and the assembly of the SJs are absolutely unknown. Given that fluctuation of intracellular Ca^2+^ levels seems to influence cell-cell communication and filament integrity (Kieninger, 2021; Walter et al., 2020), we hypothesized that cyanobacterial cell-cell communication potentially operates via the Ca^2+^-binding protein CSE.

Here, we investigated the CSE protein as a potential link between intracellular Ca^2+^ signaling and cell-cell communication. We solved the solution NMR structure of CSE in its Ca^2+^-bound state and revealed that CSE acts as Ca^2+^-buffer protein. Using cryo-electron tomography (cryo-ET), we show that CSE is not only essential for Ca^2+^ homeostasis, but also mediates the cell-cell communication via regulating the nanopores formation as Δ*cse* mutant showed a strong reduction in nanopores and SJs. This determines for the first time Ca^2+^-signaling as a novel system controlling cell-cell communication and establishes CSE as a key player regulating cyanobacterial multicellularity.

## RESULTS

### The correct folding of CSE protein depends on calcium binding

CSE is a small protein of 77 residues that is monomeric when heterologously purified as His6- tagged protein. Its sequence is highly conserved among N2-fixing multicellular cyanobacteria and contains two canonical, 12-residue EF-hand Ca^2+^-binding motifs (Fig. S1a). Accordingly, AlphaFold predicts the CSE structure with high confidence; the model shows two structurally similar Ca^2+^-binding sites, linked by an α-helix and contacting each other via a short β-bridge. It was therefore surprising that our crystallization attempts, in the presence or absence of Ca^2+^, to determine the mode of Ca^2+^-binding to CSE were unsuccessful.

Our previous results suggested that CSE undergoes substantial conformational changes upon Ca^2+^-binding (Walter et al., 2019). To explore this in more detail, we used NMR spectroscopy to determine the solution NMR structure of CSE and to investigate the dynamic of Ca^2+^-binding. For this purpose, we expressed ^13^C, ^15^N-labelled CSE and prepared an ion-free sample, using EDTA to sequester any divalent cations. The ^15^N-HSQC spectrum of this ion-free sample showed minimal peak dispersion, characteristic of an unfolded protein (Fig. S1b). Notably, two characteristic signals of folded EF-hand proteins, corresponding to glycine residues in position 6 of the EF-hand motif (Fig. S1a), were absent from this spectrum (Fig. S1b). Similarly, the 1D ^1^H spectrum displayed also little peak dispersion, with only a few defined methyl proton peaks, indicating the existence of a minimal hydrophobic core. This suggested that CSE adopts a molten globule state in the absence of Ca^2+^, consistent with our previous circular dichroism (CD) measurements (Walter et al., 2019). We therefore conclude that CSE is substantially unfolded in an ion-depleted state.

Our CD analysis indicated that both Ca^2+^ and Mg^2+^ ions can significantly induce structural changes upon binding to CSE, evident from characteristic α-helical spectra with two defined minima at 207 nm and at 222 nm (Walter et al., 2019). To further investigate the effect of divalent cations, we first titrated the ion-free CSE sample with Mg^2+^. Increasing Mg^2+^ concentration gradually improved the peak dispersion of the ^15^N-HSQC spectra, approaching that of a well-folded protein (Fig. S2). The two signals expected for the glycine residues at position 6 (Fig. S1a) of the EF-hand loops were now clearly identified at 10-11 ppm (Fig. S2a), serving as markers of a folded EF-hand. This indicates that Mg^2+^ considerably contributes to structuring the protein. However, even at 2 mM MgCl2, we were unable to record triple-resonance experiments for assignment and structure determination. This is presumably due to substantial conformational exchange associated with weaker Mg^2+^-binding, consistent with a high dissociation constant (KD) for Mg^2+^. The addition of 2 µM CaCl2 to the same sample further enhanced peak dispersion in the ^15^N-HSQC spectrum, supporting the high specificity and affinity of Ca^2+^ binding to CSE. Notably, even at high Mg^2+^ concentration (2 mM), Ca^2+^ displaced Mg^2+^ from the binding sites. Although, this sample allowed recording triple-resonance experiments, the quality of the spectra was insufficient for full sequential assignment, likely due to unfavorable exchange dynamics from competition of Mg^2+^ and Ca^2+^ for the EF-hand binding sites. Ultimately, we obtained high-quality spectra for structure determination only from a freshly prepared, ion-free sample saturated with Ca^2+^. In this Ca^2+^ saturated state, the ^15^N-HSCQ spectrum exhibited well-dispersed signals with characteristic EF-hand glycine fingerprints (Fig. 1a), consistent with a well-folded protein. Moreover, comparing the Ca^2+^- and Mg^2+^-saturated samples (Fig. S2b) revealed that several peaks shifted or disappeared in the spectrum of the Mg^2+^-saturated sample, further supporting the specificity of Ca^2+^ binding for proper CSE folding.

**Fig. 1:**
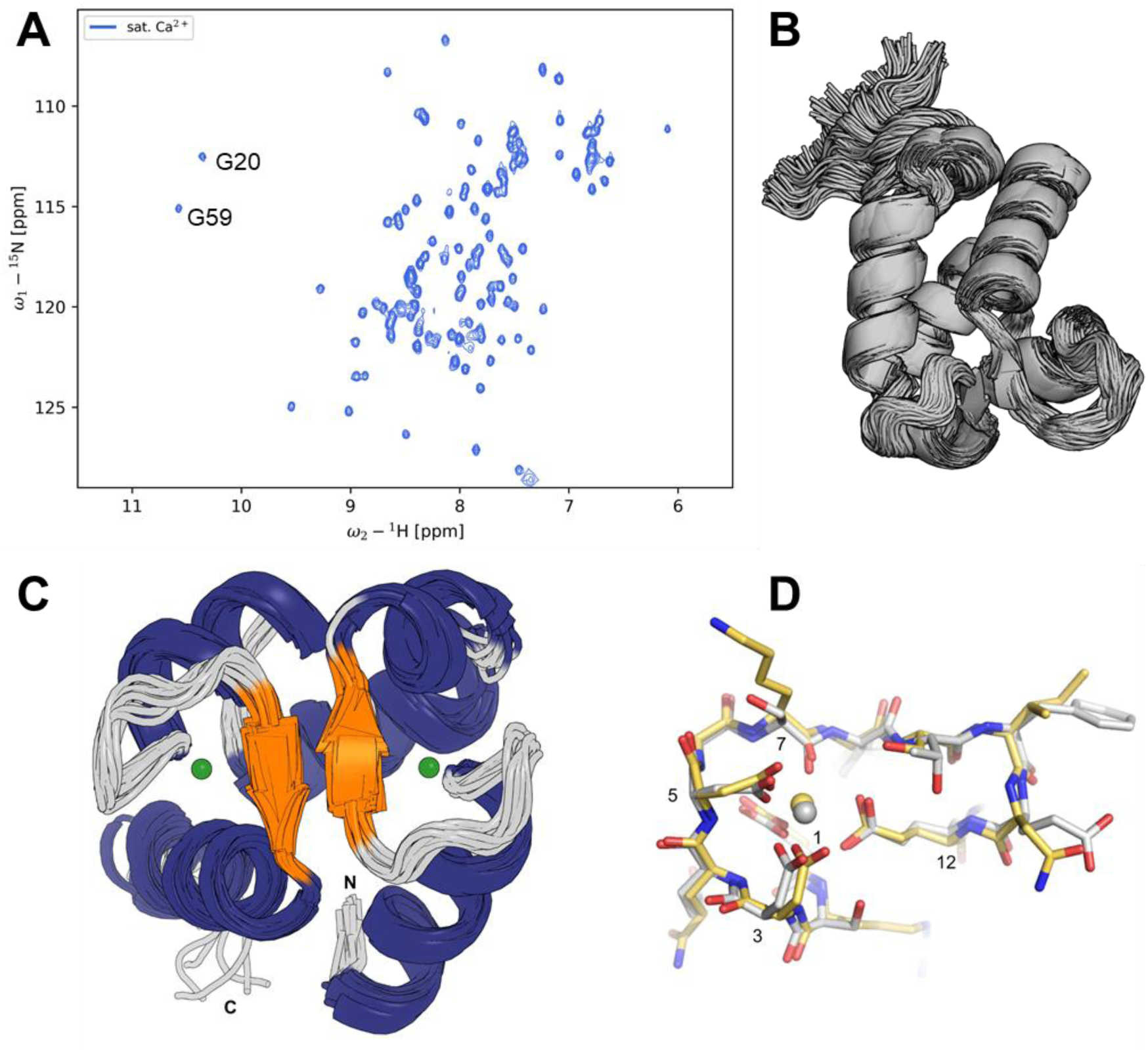
Structure of Ca^2+^-binding protein CSE. **(A)** The ^15^N-HSQC spectrum of saturated CSE:Ca^2+^ shows good peak dispersion, indicating that the protein is well folded. Two glycine residues in position 6 of the EF-hand loops (G20 and G59) are labelled. Their amide protons show an unusual downfield shift of 10-11ppm which is common among EF-hand proteins. **(B)** First 500 structural frames of solution NMR structure of CSE generated using CoMAND. **(C)** The final CoMAND ensemble for CSE colored by secondary structure. The positions of the N- and C-termini are marked. A bound Ca^2+^ ion is shown for the first model (green sphere). The backbone RMSD for superimposition of the models to the average structure is 0.66 Å. **(D)** Close-up of the CSE Ca^2+^ binding sites. The N-terminal (yellow) and C-terminal (grey) EF-hand loops are shown superimposed over backbone atoms (with RMSD 0.41 Å). Residues that coordinate Ca^2+^ are numbered according to their positions in the 12-residue EF-hand motif (corresponding to D15, D17, D19, K21, and E26 at N-terminal binding site; and D54, D56, D58, S60, and E65 at C-terminal binding site).

To determine the solution NMR structure of Ca^2+^-saturated CSE, we applied the CoMAND structure determination protocol(ElGamacy et al., 2019), which we have developed to provide an ensemble that best explains the input data and therefore reflects native-like conformational diversity. We first performed preliminary calculations without modelling of bound calcium (Fig. 1b) to confirm that residues within the EF-hand loops were well defined and positioned to make the expected pentagonal bipyramidal contacts with the ion. The CoMAND method selects frames from equilibrium molecular dynamics to best explain input NMR data from NOESY spectra. In order for these simulations to produce a realistic distribution of charges, it was necessary to model calcium ions into both sites (see material and methods). The final ensemble was selected from a pool of 2460 frames drawn from these simulations. The resulting ensemble is shown in (Fig. 1c). Structure statistics for the final ensemble are presented in (Table S1), and the final ensemble deposited under the PDB accession code 9QJW.

The Ca^2+^-bound form of CSE adopts the typical structure of a dual EF-hand protein, with two helix-loop-helix units containing the EF-hand motifs (Fig. 1c) linked by a small β-sheet. This agrees closely with the AlphaFold model, with only minor differences in backbone conformations where the final ensemble explains the data better than the starting model. The Ca^2+^-binding sites have canonical, EF-hand binding modes, with five chelating residues providing 6 ion contacts. In each case, these involve the side chains of aspartate residues in positions 1, 3 and 5 of the EF-loop, a backbone carboxyl of the residue in position 7 and a bidentate contact to the side chain of a glutamate residue in position 12. The seventh ion contact expected for pentagonal bipyramidal geometry is presumably provided by solvent. In the N-terminal site these are D15, D17, D19, K21 and E26, while at the C-terminus these are D54, D56, D58, S60 and E65 (Fig. 1d). The two loops in these structures can be superimposed with an RMSD of 0.4 Å. The CoMAND method allows a detailed analysis of the conformational diversity of these residues. In all cases, the NOESY data is well explained by a single conformation, with the R-factor expressing the match between experimental and back-calculated data approaching that of noise.

Taken together, the NMR data support the idea that CSE acts as a Ca^2+^-buffer protein, similar to other EF-hand containing proteins involved in the control of Ca^2+^-homeostasis (Domínguez et al., 2015; Yonekawa et al., 2005; Zhao et al., 2010), while it becomes unfolded in absence of divalent cations.

### CSE controls cell-cell communication in multicellular cyanobacteria

Given that Ca^2+^ seems to influence cell-cell communication (ccc) (Kieninger, 2021) and the *cse* deletion (Δ*cse*) mutant induces a strong fragmentation phenotype (Walter et al., 2020), we decided to systematically investigate the ability of CSE to control ccc in *Nostoc*. For this purpose, we used FRAP (fluorescence recovery after photobleaching) and compared the Δ*cse* knockout and *cse* overexpression (*cse* OE) mutants to wildtype cells of *Nostoc*. To test if calcium concentrations influence ccc properties, all strains were cultivated for one or two days in BG11 medium with either 0 mM, 0.25 mM (standard BG11 medium), or 1.25 mM CaCl2. Under all tested conditions, most of the cells (75-90%) of all strains showed a qualitative full recovery of the fluorescence signal and, thereby, a normal cell-cell communication behavior (Fig. S3). Also, about 5-15% of the cells exhibited a sigmoidal and/or slow increase in the fluorescence recovery, which was barely recognized for Δ*cse* cells (Fig. S3). The sigmoidal increase is characterized by a delay of up to 40-50 seconds of the fluorescence signal, then rapidly increases, like a full recovery response. Notably, neither the extracellular Ca^2+^ concentrations nor the incubation time (1 and 2 days) impact the ccc for any strain (Fig. S3).

For qualitative analysis, we measured the recovery rate of the florescent signal for the communicating cells (Fig. 2a). Under all tested conditions, the wildtype cells showed an average recovery rate of 0.074-0.090 per second. The *cse* OE strain exhibited a similar rate (0.069-0.081 per second) to the wildtype cells, while the Δ*cse* mutant clearly communicated significantly slower (0.047-0.053 per second; Fig. 2a), implying a potential change in the SJs machinery (i.e. SJs are not in the fully operative).

**Fig. 2:**
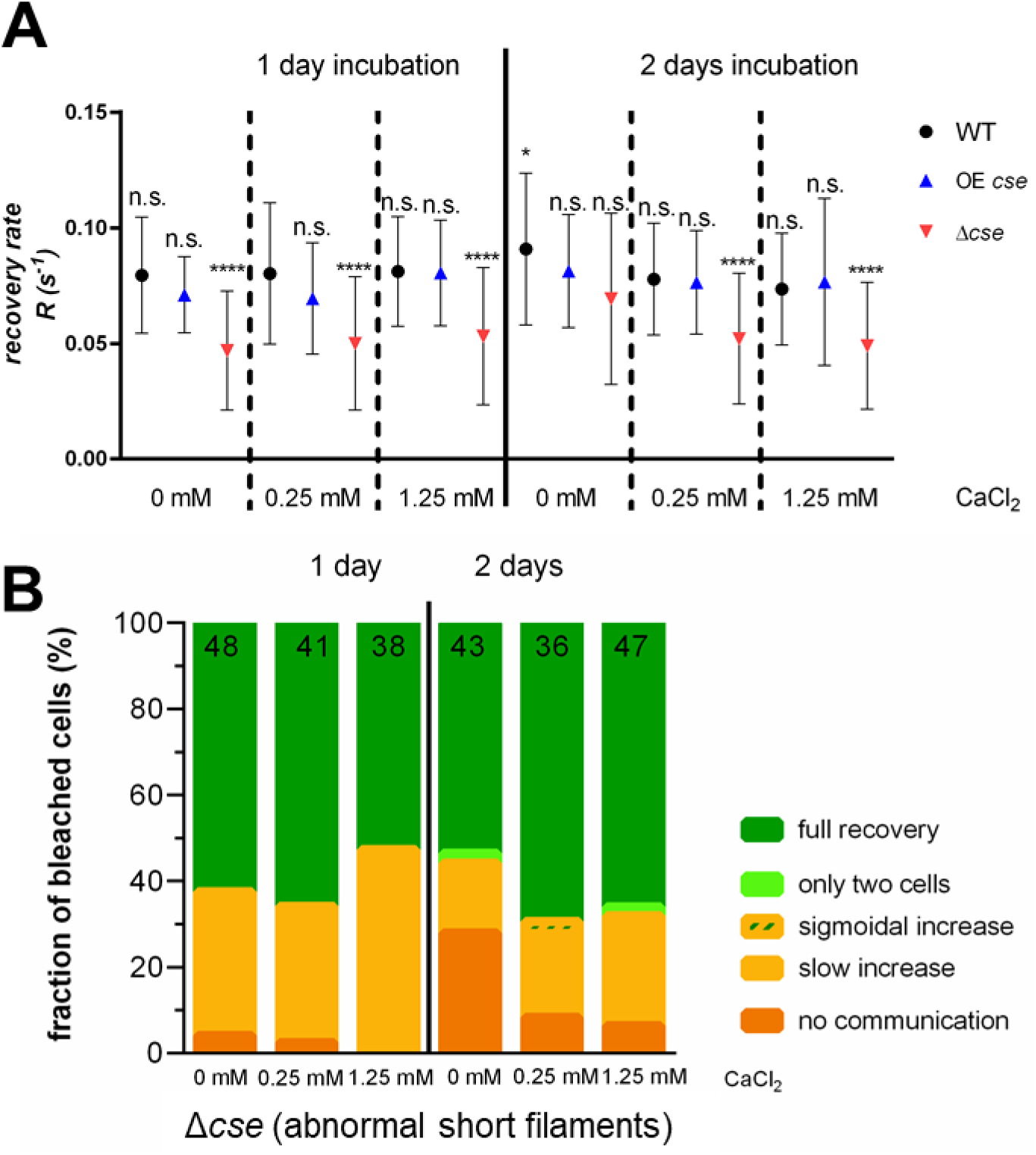
Cell-cell communication analysis via FRAP of *Nostoc* sp. PCC 7120 wildtype strain and its derived *cse* knockout (Δ*cse*) and overexpression (*cse* OE) mutants under different calcium concentrations (0 mM, 0.25mM, 1.25 mM). (A) Communication rates of *cse* mutants, calculated from FRAP analysis in (Fig. S3) and showing very slow communication for Δ*cse* mutant. The cells were incubated for one or two days with the indicated Ca^2+^ concentrations prior FRAP experiment. The results represent the mean recovery rates with standard deviation of recovering cells. Significance ***: *P* value <0.0001 in Student’s *t* test, in comparison to WT. n.s.: not significant. **(B)** FRAP analysis of the heavily fragmented short filaments of Δ*cse* mutant, showing strong impairment of cell-cell communication with 10-30% non-communicating cells.

As the Δ*cse* mutant is suffering from heavy fragmentation (Fig. 3a), we decided to additionally check the ccc within the abnormal short filaments (down to only 3 cells; Fig. 2b), which also feature unusually large cells (Fig. 3a). Here, almost 50% of the cells showed very slow ccc (slow increase) or even no ccc for 10-30% of the cells (Fig. 2b), highlighting a crucial role for *cse* in controlling cell-cell communication.

**Fig. 3:**
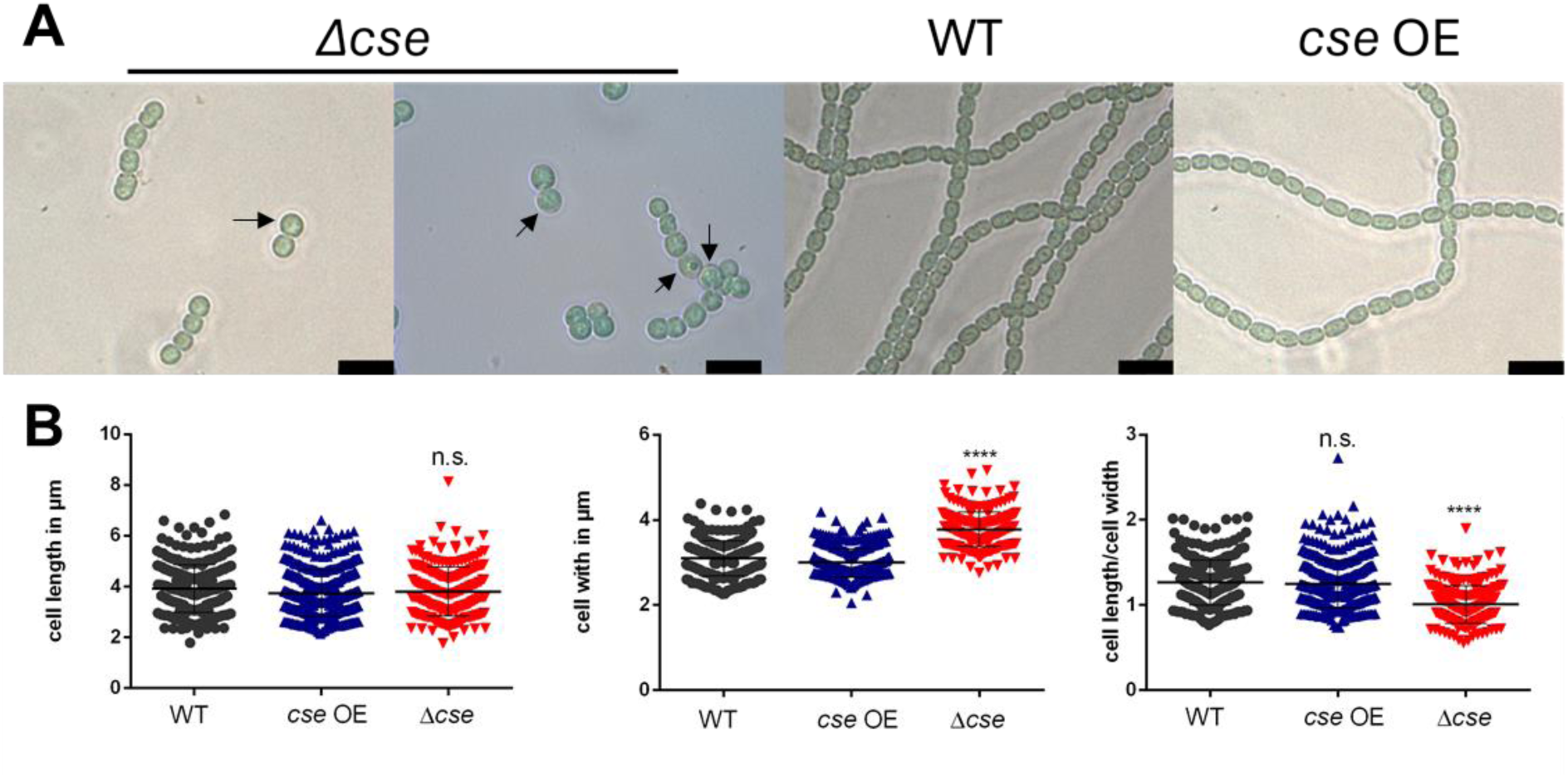
Ca^2+^-sensor protein CSE controls cell shape and filament integrity. **(A)** Light microscopy of of *Nostoc* sp. PCC 7120 wildtype strain and its derived *cse* knockout (Δ*cse*) and overexpression (*cse* OE) mutants, showing strong fragmentation phenotype of Δ*cse* mutant with irregular enlarged cells (indicated by black arrows). Bars, 10 µm. **(B)** Median values of cell length and cell width distribution of indicated strains in (panel A). Significance ****: *P* value <0.0001 in Student’s *t* test, in comparison to WT. Whiskers: standard deviation. n.s.: not significant.

In multicellular cyanobacteria, cell size homeostasis is a fundamental physical trait, which impacts on ccc and the overall cell fitness. Our observation that the short filaments of Δ*cse* mutant encounters also irregular cell shape (Fig. 3a), motivated us to systematically checked whether *cse* influences also on cell size. Interestingly, Δ*cse* cells exhibited significant cell size enlargement (particularly cell width), while *cse* OE mutant had a comparable cell size parameter to the WT cells (Fig. 3b), further supporting a vital role for Ca^2+^ signaling through CSE in multicellularity traits.

### Abundance of SJs is drastically reduced in the Δ*cse* mutant

Next, we investigated the *in situ* architecture of the ccc machinery to understand how it is altered in Δ*cse* strain. Cryo-ET of cryo-focused ion beam thinned *Nostoc* cells has proven to be a valuable tool to visualize and determine the *in situ* architecture of SJs (Weiss et al., 2019). For WT cells, we collected 22 tomograms of septa on 10 lamellae and observed SJs in 14 of them (64%) with an average of 8 SJs/tomogram (ranging from 1 to 18 SJs/tomogram; Fig. 4a and video S1). This observation agrees well with previous calculations of ∼10 SJs in a 200 nm lamella and with ∼40-80 nanopores/septum (Weiss et al., 2019). With similar parameters, we collected 14 tomograms on 10 lamellae of Δ*cse* septa and surprisingly observed SJs in only 2 of them (14%) with only 2 or 4 SJs/tomogram (Fig. 4b, Fig. S4, video S2, and video S3). The overall architecture of SJs (cap, plug and tube) in the Δ*cse* mutant appears similar to that of WT cells in the open state, however, their low abundance hindered subtomogram averaging to rule out more detailed structural changes. This striking reduction of SJs number in the Δ*cse* mutant explains the earlier results of a a slow communication (Fig. 2).

**Fig. 4:**
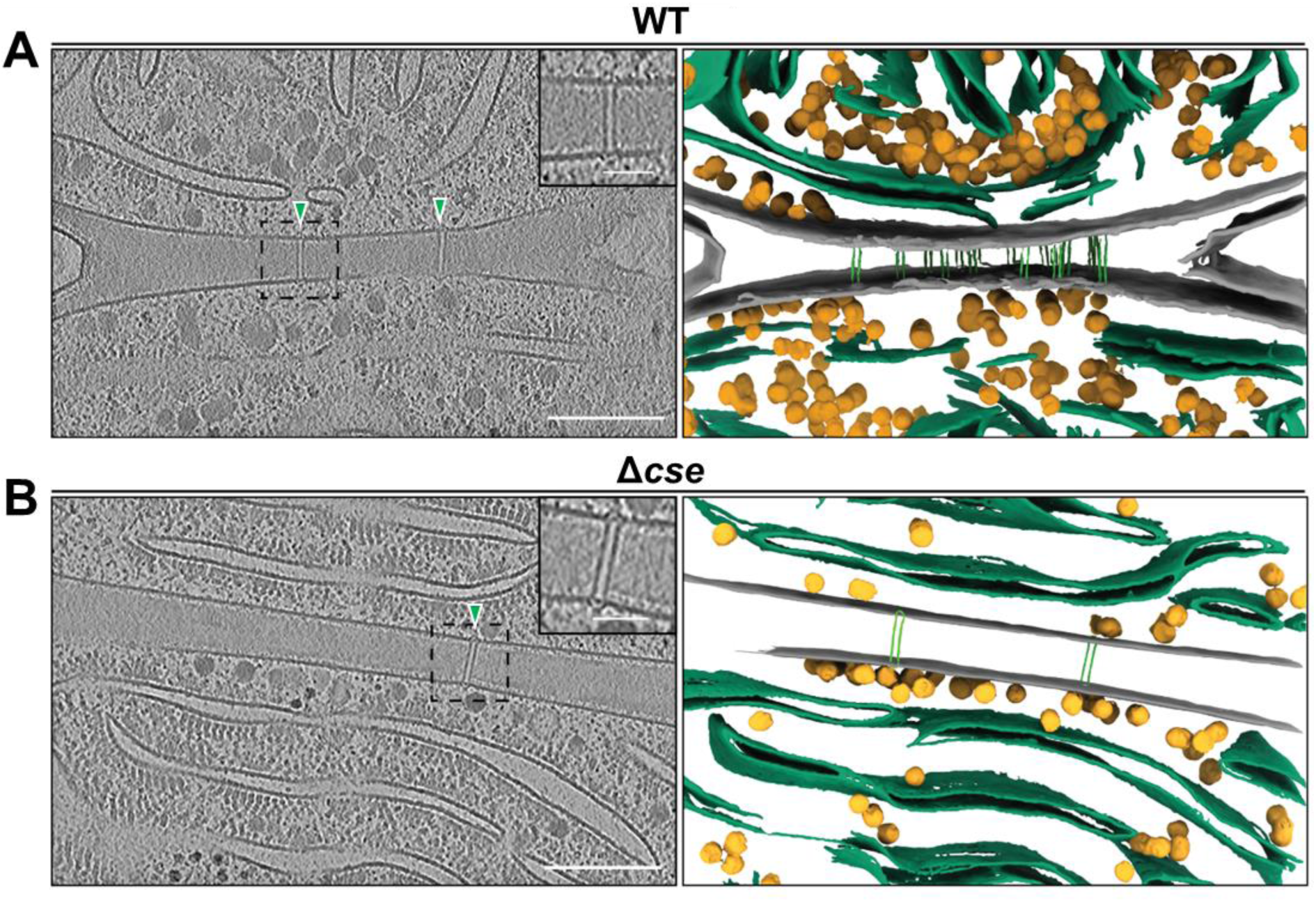
Cryo-ET reveals reduced number of SJs in Δ*cse* mutant. **(A)** Slice through a cryo-tomogram (shown is a 16 nm thick slice) of a septum of a *Nostoc* WT filament (left) and the corresponding 3D segmentation (right) shows numerous SJs (green arrowheads) per septum. In this cryo-tomogram of a ∼200 nm thick lamellae, 16 SJs were observed. A magnified view of one SJ is shown in the top right corner in the left panel. Scale bar cryo-tomogram, 200 nm. Scale bar magnified view, 50 nm. **(B)** Slice through a cryo-tomogram (shown is a 16 nm thick slice) of a septum of a *Nostoc* Δ*cse* filament (left) and the corresponding 3D segmentation (right) shows a heavily reduced number of SJs (green arrowhead) per septum. In this cryo-tomogram of a ∼200 nm thick lamellae, only 2 SJs were observed. The overall architecture of SJs in the Δ*cse* mutant appears to be similar to WT (magnified view in the top right corner of the left panel). Another example of a cryo-tomogram of a septum of a *Nostoc* Δ*cse* filament can be seen in (Fig. S4). Scale bars: 200 nm (cryo-tomogram slices) and 50 nm (magnified views).

### Molecular basis for CSE-dependent control of cell-cell communication

Previous studies have suggested a correlation between ccc, filament integrity, and proper formation of nanopores in septal PG discs (Bornikoel et al., 2017; Kieninger & Maldener 2021; Arévalo and Flores 2021; Arévalo et al., 2021; Merino-Puerto et al., 2011; Nürnberg et al., 2015). In line with this, our cryo-ET data link the highly reduced number of SJs in the Δ*cse* mutant (Fig. 4) to the strong fragmentation phenotype and the slow rate of ccc of Δ*cse* strain. This may be simply due to a reduction in the number of nanopores. Another possibility is that even if a sufficient number of nanopores are built, they might not be fully mature yet, which might hinder the proper assembly of SJs and thereby influencing on ccc negatively.

To explain the Δ*cse* phenotype mechanistically (Fig. 2 and Fig. 4) and to obtain further evidence of a functional link between CSE and ccc, we determined the efficiency of nanopore arrays formation in the Δ*cse* mutant. For this purpose, we isolated the septa septal discs of the WT and Δ*cse* strains and quantified the nanopores formation using negative staning EM (Fig. 5). It became evident that the Δ*cse* mutant possesses a huge variation in the number of nanopores compared to WT cells (Δ*cse*: 41.9 ± 34.9 nm; WT: 48,1 ± 17.8 nm) with some septa almost free from any nanopores (Fig. 5a,b), explaining our cryo-ET results. Interestingly, we additionally observed an altered septum diameter (Fig. 5c). The diameter of septal discs in WT cells agreed with the previously published data (Arévalo & Flores, 2021), while the septa of Δ*cse* cells appeared to be greatly enlarged with sizes almost double of the WT cells (Δ*cse*: 2062 ± 645 nm; WT: 1107 ± 329 nm; Fig. 5c), implying an immaturation of Δ*cse* septa. When we compared the number of nanopores to the increased septum diameter it became apparent that the Δ*cse* septal discs harbored less nanopores per µm. In this case, the increased septal area does not lead to an increased number of nanopores, further supporting the immaturation of Δ*cse* septa and explaining the slow rate of ccc (Fig. 2a). The WT exhibited a ratio of 45 nanopores per 1 µm, whereas Δ*cse* possessed only 26 nanopores per 1 µM, which is approx. 57% of the WT cells. The nanopore diameter were rather similar for both strains (Δ*cse*: 22.9 ± 5.9 nm; WT: 22.2 ± 5.0 nm). Overall, the Δ*cse* mutant exhibited a clear defect in nanopore formation, supporting a role for CSE in regulating ccc. Remarkably, several mutants which are involved in SJs (e.g. FraD) or nanopore arrays (e.g. AmiC) show fragmentation phenotype with enlargement in the septal discs and/or a reduction in the nanopores number (Kieninger et al., 2022; Arévalo and Flores 2021; Bornikoel et al., 2017), similar to that observed for Δ*cse*, further boosting an essential role for Ca^2+^ signaling via CSE in regulating ccc.

**Fig. 5.**
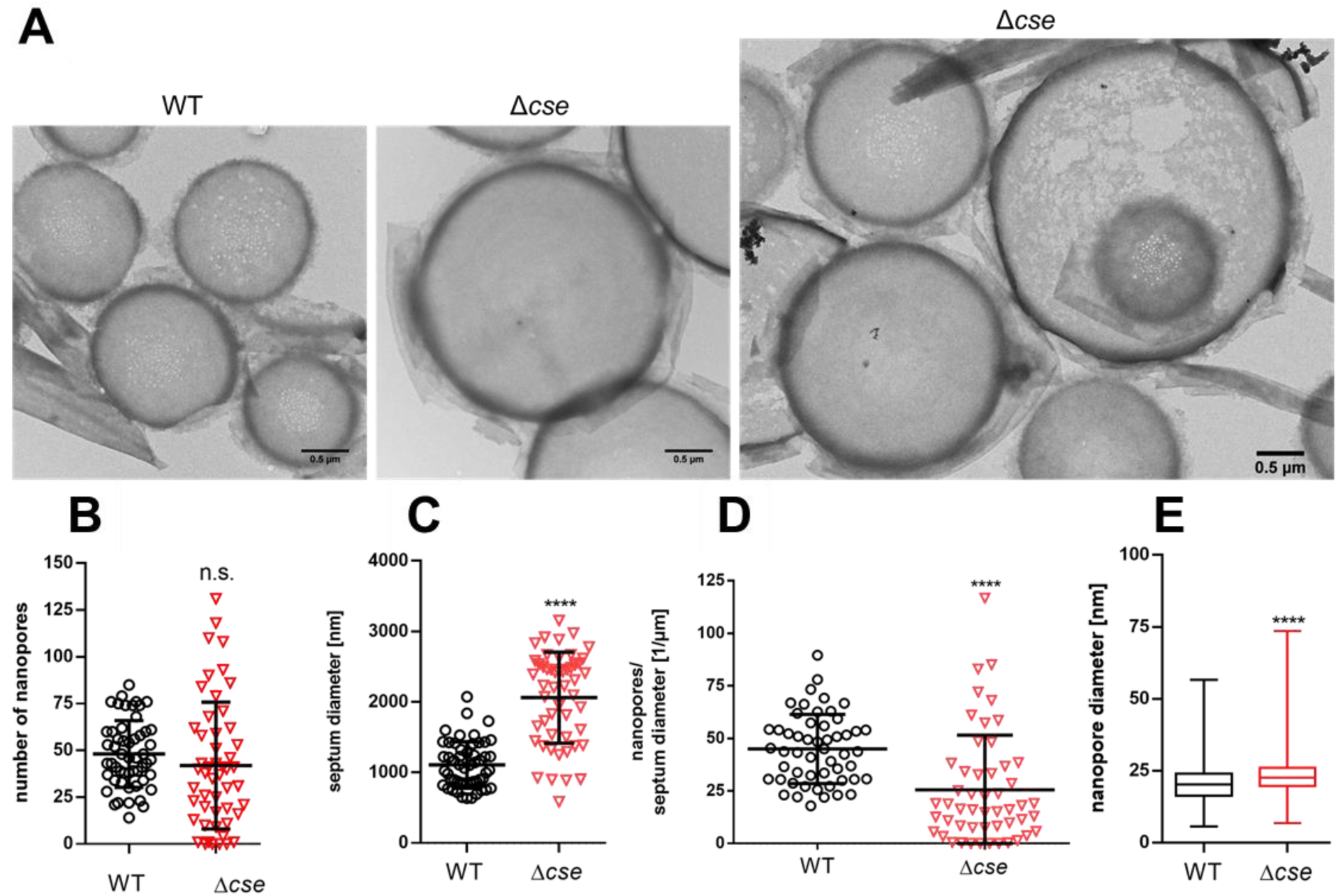
Isolated septal peptidoglycan discs of WT and Δ*cse* strains. Data of the isolated septa analysis via electron microscopy. (A) Representative septal PG discs of the WT and Δ*cse* strains. **(B)** Number of nanopores with overall mean and standard deviation (SD) of WT and Δ*cse* strains. **(C)** Septum diameter in nm with mean and SD of WT and Δ*cse* strains. (**D)** Ratio of the nanopore number per septum diameter of each data point with overall mean and SD. **(E)** Box plot of the nanopore diameter in nm of WT and Δ*cse* strains. The graphs represent the pooled data of biological triplicates.

## DISCUSSION

Filamentous cyanobacteria are one of the oldest multicellular organisms on Earth, rendering them crucial for understanding the evolutionary aspects of cell-cell communication (ccc). Cyanobacterial multicellularity includes cell differentiation and ccc for exchanging nutrients and signaling molecules between neighboring cells. We previously showed that the cyanobacterial ccc machinery is composed of nanopore arrays drilled by amidases and SJs tubes passing between the PG layers of 2 adjacent cells (Weiss et al., 2019). A strong fragmentation phenotype of cyanobacterial filaments is often associated with disrupting genes involved in controlling ccc machinery (Kieninger et al., 2022; Kieninger & Maldener 2021). As the Δ*cse* mutant is suffering from a severe fragmentation phenotype (Walter et al., 2020), we conjectured that the Ca^2+^ signaling via CSE protein may be the missing signal controlling the ccc machinery in multicellular cyanobacteria. Here, we revealed that the Ca^2+^-signaling regulates the ccc machinery, as indicated by a slow communication rate for the Δ*cse* mutant (Fig. 2). This is driven by a decrease in nanopore numbers (Fig. 4 and Fig. 5), which subsequently reduces the active SJs, rather than influencing the SJ’s architecture.

The signals, controlling cyanobacterial ccc machinery, were largely unknown. To our knowledge, CSE is the first signaling protein known to regulate ccc. This highlights Ca^2+^- signaling via CSE as a key point in cyanobacterial multicellularity, influencing not only filament integrity, heterocyst differentiation and photosynthesis (Walter et al., 2019; Walter et al., 2020), but also the ccc. The very recent discovery of an ER-plasma membrane junctions regulated by long-range Ca^2+^-signaling in neuronal dendrites (Benedetti et al., 2025) further supports our findings and suggests that the regulation of cellular junctions through Ca^2+^-signaling might be an evolutionary conserved principle between organism that diverged at least a billion years ago. This also suggests that this regulatory principle arose earlier in multicellular prokaryotes before it appeared in eukaryotes.

Our results revealed an enlargement in the Δ*cse* septa and a reduction in the nanopore numbers, similar to that observed previously for several mutants encoding for key components of the ccc machinery (e.g. Δ*fraD* and Δ*amiC*) (Weiss et al., 2019; Kieninger et al., 2022; Arévalo and Flores 2021; Bornikoel et al., 2017). Large septal discs are usually observed in young, immature divided cells, explaining the few nanopores observed. Given the strong fragmentation phenotype of Δ*cse* mutant and its negative influence on the nanopore arrays and septa, it’s possible that CSE is additionally regulating the cell division or the cell wall maturation via amidases (AmiC), which are involved in drilling and maturation of nanopores. *Nostoc* possesses three amidases, while AmiC1 and AmiC2 are involved in nanopores formation, the AmiC3 regulates the nanopore size (Lehner et al., 2013; Bornikoel et al., 2017; Zheng et al., 2017). It’s largely unknown how the cell wall amidases are regulated, therefore it’s possible that CSE interacts directly with one or more of AmiCs, similar to LytM, which interact with AmiC1 and enhances its activity (Bornikoel et al., 2018). As CSE acts as Ca^2+^- buffer protein and Ca^2+^ is known to operate more broadly in metabolic signaling, it is also possible that free Ca^2+^ ions directly control the amidases activity, induce conformational changes on the SJs components, or influence the levels of proteins transcriptionally or translationally. Notably, external Ca^2+^ supplementation had no influence on the communication rate, consistent with the tight control of free cytoplasmic Ca^2+^ ions (at nM levels) through Ca^2+^ pumps and Ca^2+^-buffer proteins (Mantovani et al., 2023).

Our structural analysis supports the notion that CSE is a Ca^2+^-buffer protein. In absence of divalent ions, CSE is substantially unfolded, however both Ca^2+^ (and to some extent Mg^2+^) ions can induce folding, supporting the high specificity of CSE towards Ca^2+^-binding. While Ca^2+^ concentrations are strictly controlled at nM levels, Mg^2+^ is one of the most abundant metal ions in the cytoplasm. CSE therefore likely exists in a partially folded ‘waiting’ state at low Ca^2+^ concentrations, and the buffering capacity of CSE is likely controlled by the relatively high affinity for Ca^2+^ compared to Mg^2+^ ions (Walter et al., 2019). CSE possesses two EF-hand Ca^2+^-binding motifs responsible for sequestering free cytoplasmic Ca^2+^ ions. Similarly, several EF-hand Ca^2+^-binding proteins were shown to act as Ca^2+^-buffer proteins, further supporting our finding. For example, *Pseudomonas aeruginosa* encodes a EF-hand Ca^2+^-buffer protein, EfhP (Sarkisova et al., 2014), while the filamentous actinobacterium *Streptomyces coelicolor* harbors four EF-hand Ca^2+^-binding proteins (CabA, CabB, CabC and CabD), suggesting a role for Ca^2+^ EF-hand proteins in multicellularity (Domínguez et al., 2015; Wang et al., 2008; Yonekawa et al., 2005; Zhao et al., 2010). Filamentous cyanobacteria possess another conserved Ca^2+^-binding protein, known as CcbP (Zhao et al., 2005; Hu et al., 2011). In contrast to CSE, CcbP seems to be implicated mainly in heterocyst differentiation and Ca^2+^-binding does not induce strong conformational changes on its structure. CcbP captures free Ca^2+^ ions via a noncanonical high affinity binding site with an α-turn-β region and a weak site with an EF-hand-like motif.

Though only two Ca^2+^-binding proteins are known and analyzed in *Nostoc*, there are vast majority of putative EF-hand proteins in cyanobacteria. CSE protein appears to be very important for multicellularity, as - with a few exceptions - it is only found in filamentous cyanobacteria and plays important roles in regulating filament integrity, heterocyst differentiation and ccc machinery (Walter et al., 2019).

## Material and Methods

### NMR Spectroscopy

For NMR structure determination, N-terminally His6-tagged CSE protein was expressed in minimal medium containing ^13^C-glucose and ^15^N-ammonium chloride at 37 °C by induction with IPTG. The purification process involved Ni-NTA chromatography followed by size exclusion chromatography using an Äkta system as described previously (Walter et al., 2019). The protein was eluted in a 20 mM Tris-HCl buffer (pH 7.9) containing 100 mM NaCl.

NMR experiments were conducted at 293 K on 600 MHz Bruker Avance-III and 800 MHz Bruker Avance-III spectrometers. Backbone sequential assignments were made using standard triple-resonance experiments. Note that these were only possible a freshly prepared ion-free sample that was subsequently saturated with 2 mM CaCl2. As the CoMAND structure determination protocol employs a ^13^C-edited 3D-CNH-NOESY, the aim of the side chain assignment process was to achieve the most complete ^13^C assignment possible. To this end we used dedicated TOCSY-based experiments for both aliphatic and aromatic sidechain assignments.

Structure determination of Ca^2+^-saturated CSE was performed using the CoMAND protocol (ElGamacy et al., 2019). The input data for this protocol was derived from a 3D-CNH-NOESY spectrum (Diercks et al., 1999) obtained at 800 MHz (mixing time 120 μs). The processed 3D matrix was first broken into a set of 1D vectors perpendicular to assigned ^15^N-HSQC positions, such that each contains peaks to one ^15^N-bound proton. The CoMAND method is based on selecting conformers from a pool in order to best explain to these experimental vectors (strips). This is judged by a quantitative R-factor reflecting the match between experimental and back-calculated data. As each strip is associated with a single ^15^N-bound proton, the CoMAND R-factor is calculated on a per-residue basis.

The starting point for the CoMAND protocol is a model of the folded protein. For this purpose, we chose the AlphaFold2.0 model, which has very high confidence scores on the level of the global fold. In order to sample conformational diversity, we used standard simulated annealing protocols in XPLOR-NIH (Schwieters et al., 2006; Schwieters et al., 2003), using generic distance and dihedral angle restraints generated from the starting model. These runs have the advantage of efficient sampling of conformational space for loops and unstructured regions of the protein. These simulations were run with Ca^2+^ bound via geometric restraints based on the canonical EF-hand binding mode and pentagonal bipyramidal Ca^2+^ geometry. A total of 50 structures were calculated, from which we selected the best 8 models, based on their overall R-factor. The selected structures were then used as individual starting points for unrestrained molecular dynamics simulation runs in explicit solvent using NAMD 2.12 (Phillips et al., 2020) with an underlying CHARMM36 forcefield.

For the first unrestrained MD runs we sampled a total of 4000 frames from which we again selected based on R-factor. In this case we selected at the per-residue level; i.e. a set of MD frames were selected to optimize the R-factors for each residue for which strips were available. This optimization provides detailed information on the conformational distributions for each residue. In order to investigate the conformational diversity of residues expected to contact the calcium ion, we carried out these simulations without bound calcium. It should be noted that conformations consistent with calcium binding are under-sampled in such simulations, due to unfavorable charge distributions. Nevertheless, sufficient information on the distributions was obtained for each residue to confirm the canonical calcium binding mode. The sets of frames collated for each residue were used to compile distance and dihedral restraints that effectively replace those based on the original model. These were used in a second XPLOR-NIH run to generate a new set of 400 structures.

At this stage, it is expected that optimization based on single residues should explain the experimental data very well - in most cases approaching the boundary represented by experimental noise and the accuracy of back-calculation. For some residues, this was not the case, suggesting that the conformations present in the original AlphaFold2.0 model were not sufficient to explain the data. These included two backbone polymorphisms that are typically not well predicted by static AlphaFold2.0 models. In the case we employed exhaustive grid searches to identify any upsampled conformers and built models containing the alternate conformation in XPLOR-NIH using appropriate distance and dihedral restraints. A set of 12 structures respecting these polymorphisms was compiled as the starting point for a final round of MD sampling, with each used to seed a short trajectory at 310 K in explicit solvent, in this case conducted with calcium modelled into the binding sites. We selected the final ensemble from this pool of 2460 frames.

### Cyanobacterial strains and verification of segregation of Δ*cse*

*Nostoc* sp. PCC 7120 was used as wildtype (WT) strain. The Δ*cse* strain was gifted from Peter Gollan laboratory (University of Turku, Finland). In this strain the gene coding for CSE, *asr1131*, was replaced by a Neomycin resistance cassette (Walter et al., 2020). The overexpression strain was generated as described previously (Walter et al., 2019). *Nostoc* sp. PCC 7120 and the *cse* mutants were cultivated in blue green media with sodium nitrate (BG11; shown is table S2) (Rippka et al., 1979). Liquid cultures were grown under agitation at 100 rpm and plate cultures were grown on solidified BG11 with 1.5% (w/v) Bacto agar (Becton Dickinson) at 28 °C and constant illumination at 30-40 µE m^-2^ s^-1^. The full segregation of the Δ*cse* strain was tested by colony PCR (Fig. S5) using high-fidelity polymerase Q5 (NEB) and different primer sets (Table S3).

### Fluorescence recovery after photobleaching (FRAP) analysis for measuring cell-cell communication

The fluorescence dye calcein was used to assess cell-cell communication as stated earlier (Merino-Puerto et al., 2011; Mullineaux et al., 2008). The method was further modified according to (Weiss et al., 2019) to test the cell-cell communication under different calcium chloride concentrations (0 mM, 0.25 mM, 1.25 mM CaCl2). The day before FRAP measurements cells from liquid preculture (OD750 0.6-1, BG11) were washed three times in 1 ml BG11 medium with respective calcium chloride concentration and spread on 1.5 % (w/v) agar plates with the respective calcium chloride concentration. On the day of measurement, the cells were washed three times in medium and resuspended to an OD750=2.4 in 250 µl. The fluorescence dye calcein-AM (stock 1 mg·mL^-1^ in DMSO, Thermo Fisher) (4-6 µl) was added. The samples were incubated for 90 min at 28°C in the dark under constant gentle agitation and washed three times at 1700 *xg* for 2 min with medium afterwards. The cells were resuspended in 500 µl BG11 medium and further incubated for 90 min. For FRAP, 10 µL of each sample were spotted onto a BG11 agar plate with respective calcium chloride concentration. The dried samples were studied with a Zeiss LSM 800 confocal microscope. For normal cells the middle cell of a filament consisting of at least five equally dyed cells was bleached. For the Δ*cse* strain shorter filaments (3 cells) with deviating cell size were assessed as well. Before bleaching five pictures were taken. Pictures at 1.1 s interval were taken for 60- 100 s to record the fluorescence recovery in the bleached cells. According to (Kieninger, 2021; Weiss et al., 2019) the programs GraphPad Prism and ImageJ were used to analyze the relative fluorescence in the bleached cells over time.

### Septa isolation and transmission electron microscopy

To assess the properties of the septal disc, the septa were isolated according to (Kieninger, 2021; Kühner et al., 2014). Only filtered (0.22 µm) solutions were used. Cells from exponentially growing liquid cultures of *Nostoc* strains were spread on BG11 plates and incubated for 3 days under standard growth conditions. One third to one half of the cell material were scratched and resuspended in 1 ml ultrapure water and centrifuged at 10 000 x*g* for 5 min. For fragmentation and mechanical disruption of the membranes the cell pellets were resuspended in 700 μl 0.1 M Tris buffer pH 6.8 and sonicated (duty care=50, output control=1) for 2 min. Subsequently 300 µl of 10% SDS was added for denaturation. To melt the membranes the samples were incubated at 99°C and 300 rpm for 25 min. After centrifugation at 10 000 x*g* for 5 min the pellet was washed twice with 1 ml ultrapure water. The peptidoglycan sacculi were mechanically degraded by incubation in a sonification water bath for 30 min. The samples were again centrifuged at 10 000 x*g* for 5 min. The pellet was resuspended in 1.5 ml 50 mM Na3PO4 solution (pH 6.8) and incubated for 3 h with 15 μl α-Chymotrypsin (300 µg) added for enzymatic proteolysis. Further 15 μl α-Chymotrypsin (300 µg) were added and the reaction mix was incubated overnight. Via incubation at 99°C for 3 min the enzyme was inactivated, and the mix was centrifuged. The quality of the sample was assessed in the fluorescence microscope by the addition of 1% (v/v) fluorescent vancomycin, (Vancomycin, BODIPY FL conjugate, 100 μL/mL in DMSO, Invitrogen). If required additional sonication steps for 30-40 s followed. After centrifugation the pellet was resuspended to a final volume of 300 – 500 µl from which 10 µl were applied to UV-irradiated formvar/carbon film-coated copper grids. After 30-60 min incubation the grids were washed with ultrapure water and dried with filter paper before incubation with uranyl acetate (1% in ultrapure water) for 1 min. Dried grids were assessed with transmission electron microscopy using a Philips Tecnai10 electron microscope at 80 kV connected to a Rio Camera (Gatan) for imaging. Transmission electron micrographs were analyzed as described in (Kieninger, 2021).

### Plunge freezing of *Nostoc* filaments

Prior to plunge freezing, *Nostoc* filaments grown on BG11 plates were resuspended in liquid medium to obtain a dense cell suspension. 3.5 μl of this cell suspension was applied to glow-discharged copper EM grids (R2/2, Quantifoil) and automatically blotted from the back for 12- 14 s using a Vitrobot plunge-freezing robot (Thermo Fisher), in which one blotting paper was replaced with a Teflon sheet (Weiss et al., 2017). The sample was subsequently vitrified by plunging into liquid ethane-propane (Tivol et al., 2008), and frozen grids were further stored in liquid nitrogen.

### Cryo-Focused Ion Beam milling

Electron-transparent lamellae through plunge-frozen *Nostoc* filaments were prepared using the SerialFIB software package (Klumpe et al., 2021) on a Zeiss Crossbeam 550 FIB-SEM dual-beam instrument (Carl Zeiss Microscopy, Oberkochen). EM grids were clipped into FIB milling autoloader-grids (ThermoFisher Scientific, Waltham, MA) and mounted onto a 40° pre-tilted grid holder (Medeiros et al., 2018) (Leica Microsystems GmbH, Vienna, Austria) using a VCM loading station (Leica Microsystems GmbH, Vienna, Austria). Cryo-grid transfer was performed with a VCT500 cryo-transfer system (Leica Microsystems GmbH, Vienna, Austria). Grids were sputter-coated with a ∼8 nm thick layer of tungsten using an ACE600 cryo-sputter coater (Leica Microsystems GmbH, Vienna, Austria) and afterward transferred into the dual beam instrument equipped with a copper-band cooled mechanical cryo-stage (Leica Microsystems GmbH, Vienna, Austria). An organometallic platinum precursor layer was deposited onto each grid via the gas injection system, and the SEM (3–5 kV, 58 pA) was used to identify suitable milling targets. Milling patterns were placed and executed via the SerialFIB software package, aiming for a final lamellae thickness of ∼200 nm by applying gradually reducing currents for rough milling (700, 300, and 100 pA) and for polishing (50 pA). Grids with final lamellae were unloaded and stored in liquid nitrogen until cryoET imaging.

### Cryo-electron tomography and tomogram reconstruction

Tilt-series of FIB-milled lamellae were collected on a Titan Krios 300kV FEG transmission electron microscope (Thermo Fisher) equipped with a K3 direct electron detector (Gatan) combined with a BioContinuum imaging filter (slit width 20eV). All data were acquired using the SerialEM (Mastronarde, 2003, 2005) and PACEtomo (Eisenstein et al., 2023) software packages. Tilt-series were recorded with a dose-symmetric tilt-scheme covering 120° of angular range, with a pixel size of 2.68 Å at the specimen level, at -6 μm defocus, and with a cumulative electron dose of ∼140e− Å−2.

The acquired images were further processed with the IMOD package (Kremer et al., 1996; Mastronarde, 2008). After motion-correction with ‘alignframes’, the tilt-series were reconstructed into 4x binned tomograms using weighted back-projection. Cryo-tomograms were furthermore filtered using IsoNet (Liu et al., 2022) to improve contrast for visualization. Segmentations were created using Dragonfly (Heebner et al., 2022) and subsequently imported into ChimeraX (Pettersen et al., 2021), Gaussian filtered and processed with the surface smoothing function. All segmentations were visualized using ChimeraX.

## Data availability

All data supporting the findings of this study are available within the paper. The NMR data, atomic coordinates, and structure factors have been deposited in the Protein Data Bank, www.wwpdb.org (PDB ID codes: 9QJW). The cryo-electron tomograms shown in figures were uploaded to the Electron Microscopy Data Bank (EMDB) with accession numbers EMD-52774 – EMD-52776.

## Acknowledgements

This work was supported by the German research foundation (DFG) within the priority program SPP2389 (SE 3449/1-1) and Emmy Noether program (SE 3449/3-1) to KAS, and by institutional funds of the Max Planck Society. KAS also gratefully acknowledges the infrastructural support and funding by the collaborative research center SFB1535 (MibiNet) and the Excellence Strategy of the German Federal and Baden-Württemberg State Governments (Projektförderung: PRO-SELIM-2022- 14). We are grateful to Jörg Scholl, Ana Janović, Claudia Menzel, and Heinz Grenzendorf (IMIT, Tübingen University) for the excellent assistance. Furthermore, we would like to acknowledge Martin Pilhofer (ETH Zürich), Karl Forchhammer and Iris Maldener (Tübingen University) for continued support and constructive discussions. Also, we would like to thank Sven Klumpe, Jürgen Plitzko (Max-Planch Institute for Biochemistry, Martinsried), and Roland Salzer (Zeiss) for developing and implementing SerialFIB for the Zeiss Crossbeam, and Peter Gollan (Turku University) for sharing materials and strains with us. ScopeM is acknowledged for cryoEM instrument access at ETH Zürich.

## Author contributions

TM and KAS conceptualized, initiated and designed the research. KAS supervised the study. TM performed all physiological experiments. KR and GW performed and analyzed the cryo-ET data. KAS initiated the project, acquired 3^rd^ party funding and prepared samples for the NMR experiments. FMK and MC performed the NMR experiments and solved the CSE structure. TM and KAS wrote the initial draft of the manuscript. All authors edited and approved the final manuscript.

---

**Fig. S1:**
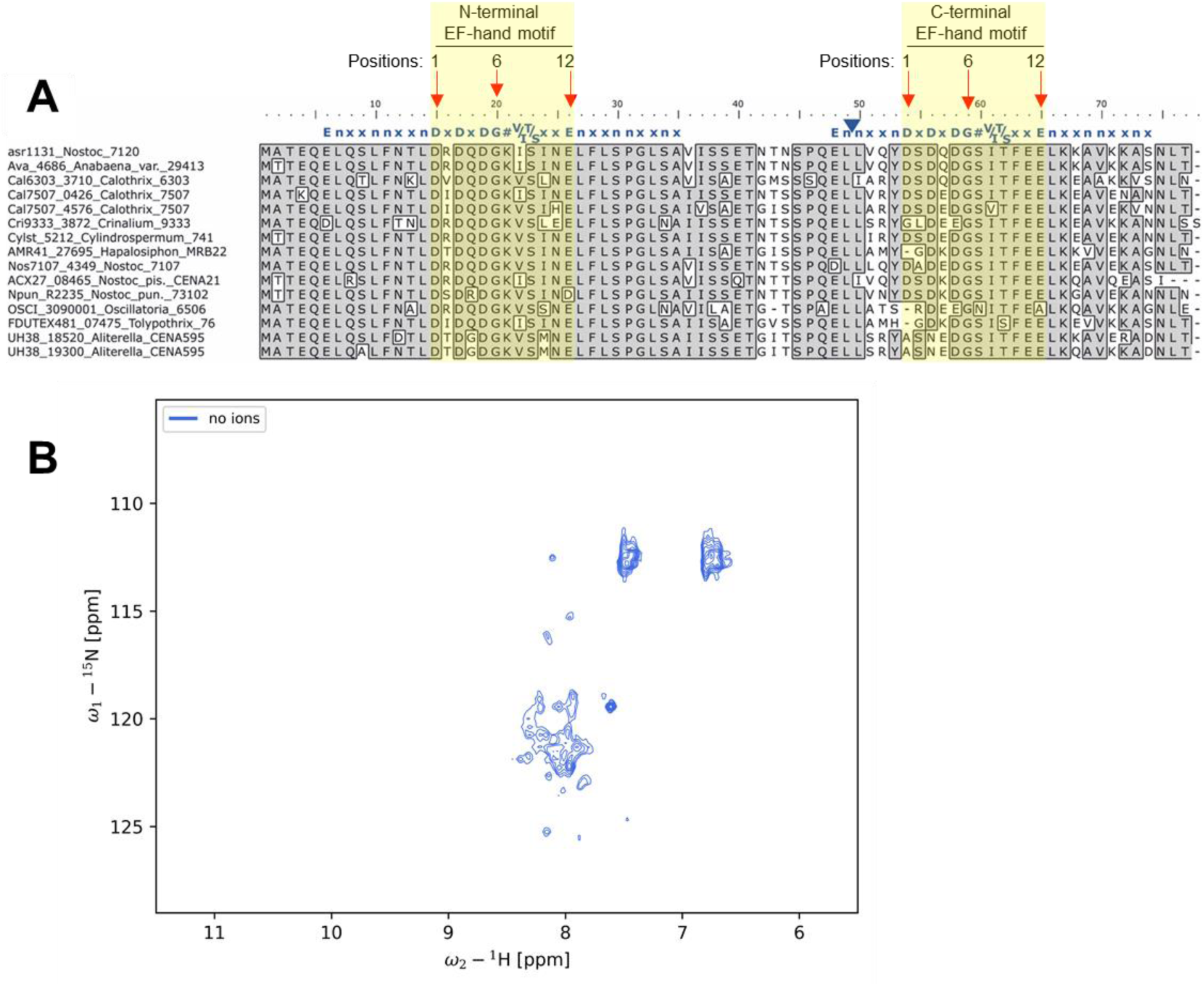
(A) Multiple sequence alignment shows conservation of CSE proteins among N2-fixing multicellular cyanobacteria with 2 characteristic EF-hand Ca^2+^-binding motifs (positions 1-12; yellow boxes). **(B)** ^15^N-HSQC spectrum of the ion-free CSE protein (i.e. without Mg^2+^ or Ca^2+^ ions).

**Fig. S2:**
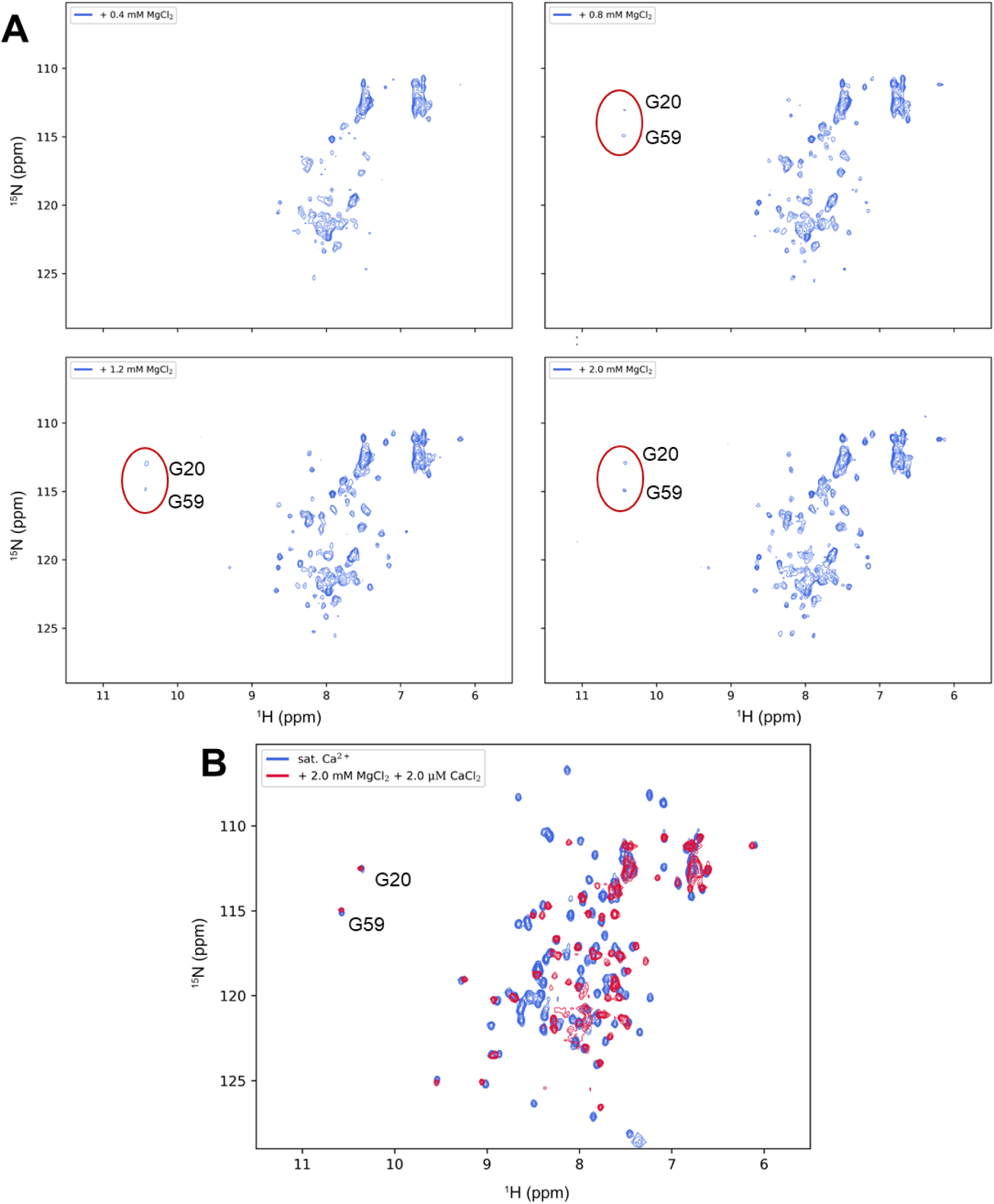
^15^N-HSQC spectra of CSE with Mg^2+^ and/or Ca^2+^. **(A)** ^15^N-HSQC spectra of CSE samples titrated with increasing concentrations of MgCl2 (0.4, 0.8, 1.2 and 2.0 mM). Peaks corresponding to the characteristic glycine residues in position 6 of the EF-hand loops (G20 and G59) are labelled. **(B)** An overlay of the ^15^N-HSQC spectra of CSE saturated with Ca^2+^ (blue) and CSE titrated with 2 mM Mg^2+^ (red). The considerable differences in both peak positions and overall dispersion indicate that Ca^2+^ efficiently competes with Mg^2+^ for binding sites.

**Fig. S3:**
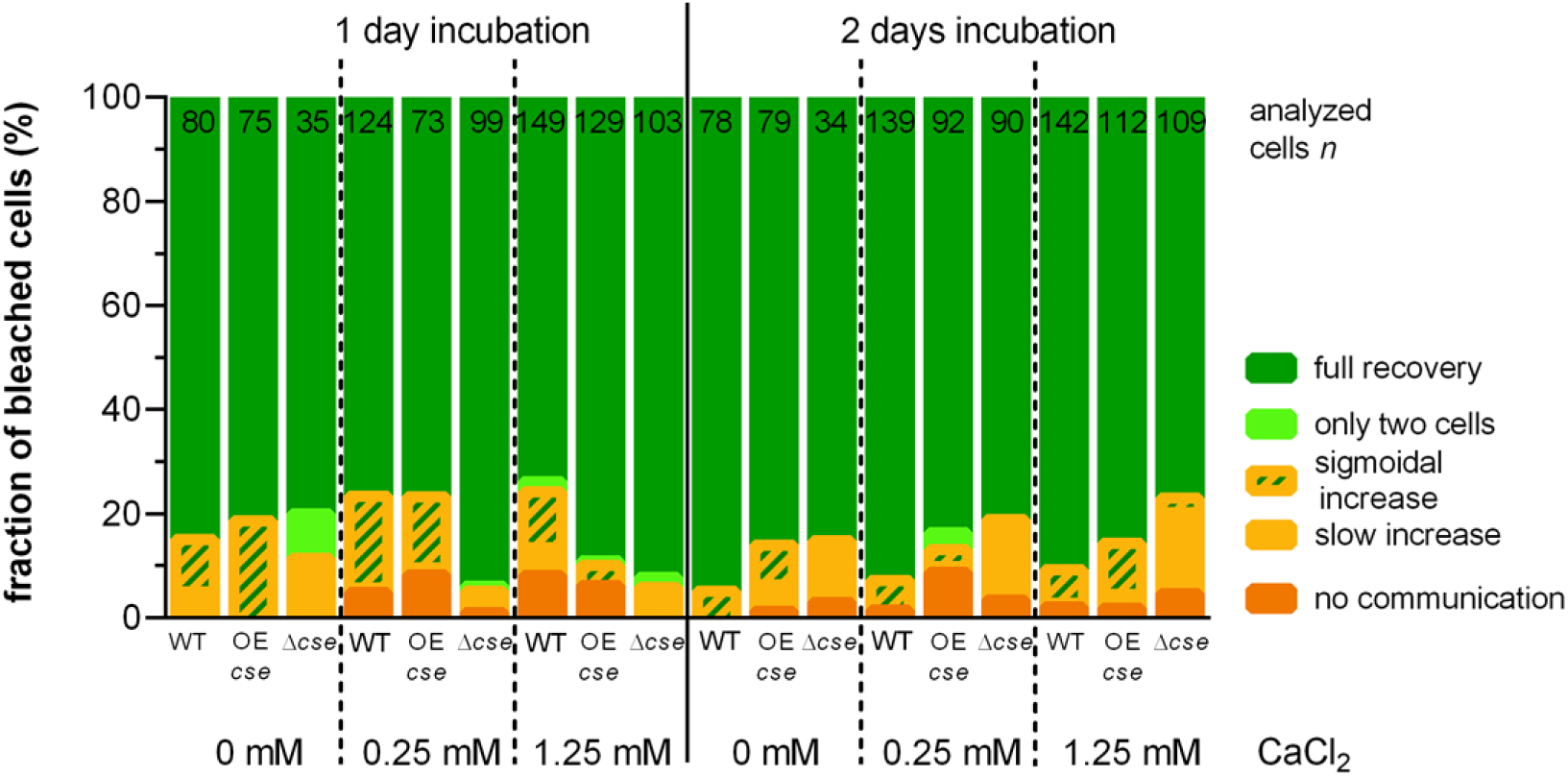
Cell-cell communication analysis via FRAP of *Nostoc* sp. PCC 7120 wildtype strain and its derived *cse* knockout (Δ*cse*) and overexpression (*cse* OE) mutants under different calcium concentrations (0 mM, 0.25mM, 1.25 mM). The cells were incubated for one or two days with the indicated Ca^2+^ concentrations prior FRAP experiment. Pooled data is shown from at least biological quadruplets, except for Δ*cse* mutant at 0 mM CaCl2 (with biological duplicates).

**Fig. S4:**
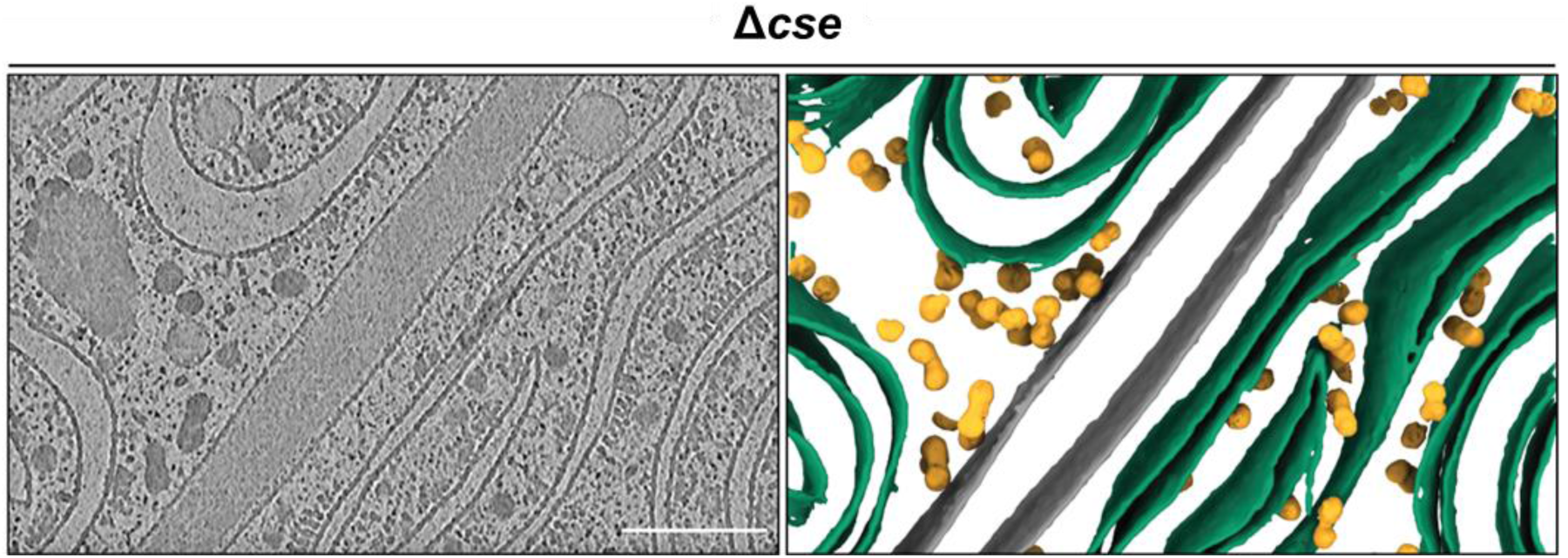
Cryo-ET reveals highly reduced number of SJs in Δ*cse* mutant. Slice through a cryo- tomogram (shown is a 16 nm thick slice) of a septum of a *Nostoc* Δ*cse* filament (left) and the corresponding 3D segmentation (right) shows a heavily reduced number of SJs per septum. In this cryo- tomogram of a ∼200 nm thick lamellae, no SJs were observed. In comparison, ∼8 SJs/tomogram can be observed in WT filaments. Scale bars: 200 nm (cryo-tomogram slice) and 50 nm (magnified view).

**Fig. S5:**
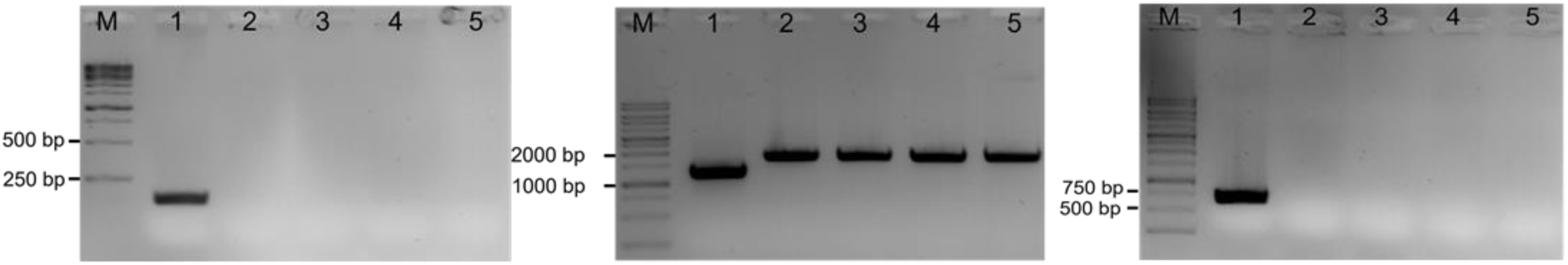
Genotypic characterization of *Δcse* knockout mutant. PCR using different sets of gene- specific primers (see Supplementary Table S3) showing complete segregation of *Δcse* knockout mutant with insertion of neomycin-resistance-cassette. Lanes: 1 (WT *Nostoc* cells), while 2-5 (Δ*cse* cells).

**Table S1:**
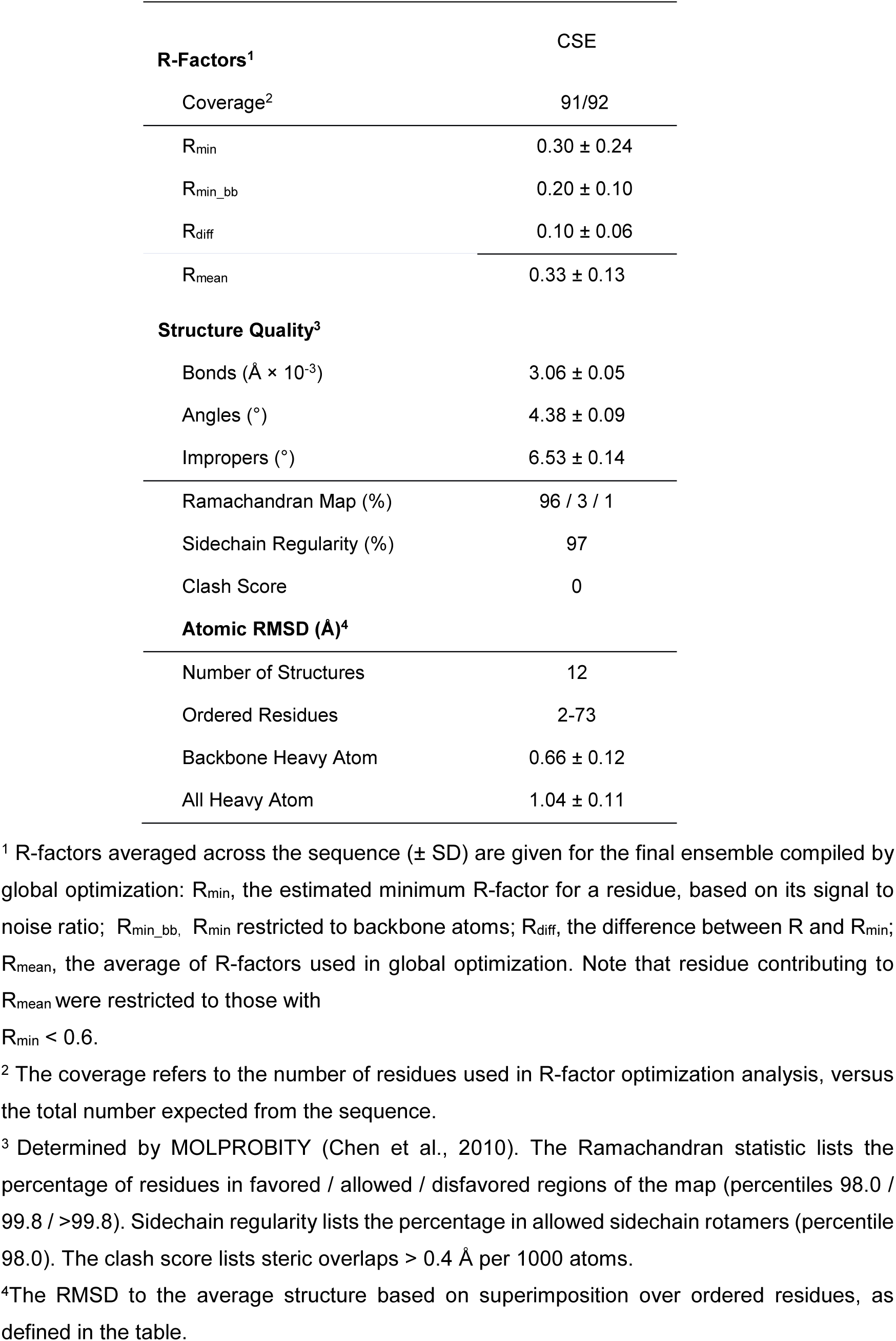
CoMAND ensemble structure statistics of CSE protein, related to Fig. 1.

**Table S2:** Composition of BG 11 media used for *Nostoc* cultivation.

| component | μM | component | μM |
| --- | --- | --- | --- |
| MgSO<sub>4</sub> x 7H<sub>2</sub>O | 304 | H<sub>3</sub>Bo<sub>3</sub> | 46.3 |
| CaCl<sub>2</sub> x 2H<sub>2</sub>O | 245 | MnCl<sub>2</sub> x 4 H<sub>2</sub>O | 9.2 |
| K<sub>2</sub>HPO<sub>4</sub> | 175 | ZnSO<sub>4</sub> x 7 H<sub>2</sub>O | 0.77 |
| Na<sub>2</sub>CO<sub>3</sub> | 180 | Na<sub>2</sub>MoO<sub>4</sub> x 2 H<sub>2</sub>O | 1.6 |
| C<sub>6</sub>H<sub>8</sub>O<sub>7</sub> | 31 | CuSO<sub>4</sub> x 5 H<sub>2</sub>O | 0.32 |
| FeC<sub>6</sub>H<sub>5</sub>O<sub>7</sub> | 24 | CoCl<sub>2</sub> x 6 H<sub>2</sub>O | 0,2 |
| Na<sub>2</sub>EDTA x 2 H<sub>2</sub>O | 3.4 | NaNO<sub>3</sub> | 17.6 x10<sup>3</sup> |

**Table S3:**
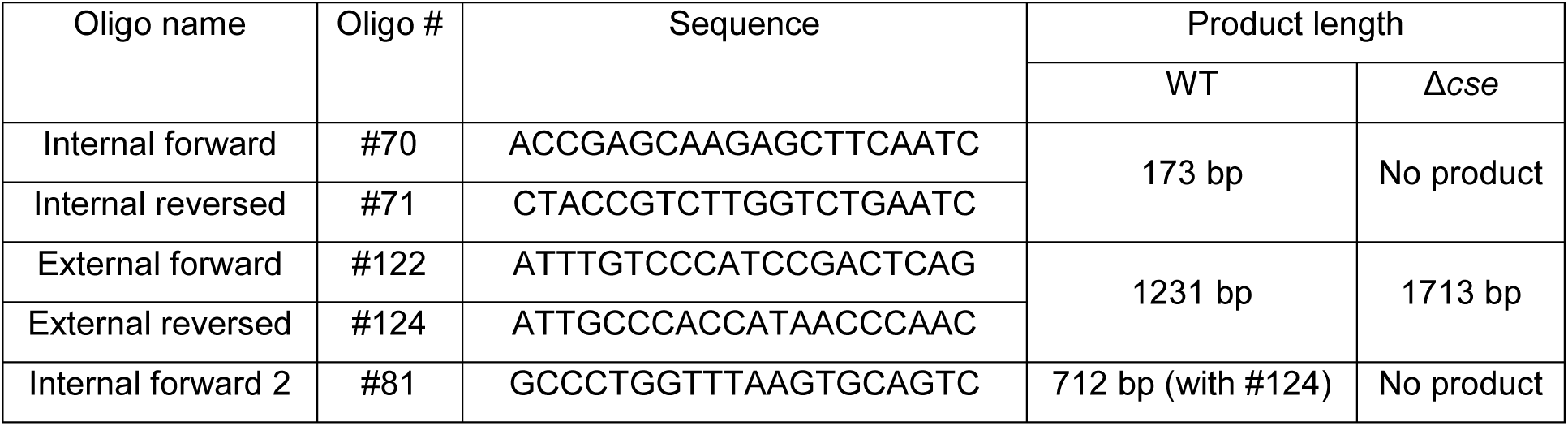
Primers used in this study. Different oligonucleotides, which bind inside or up- and downstream of *cse* gene.

**Video S1.** Cryo-tomogram of a septal region of wildtype *Nostoc* followed by segmentation, related to (Fig. 4a) and shows normal abundance of septal junctions. Septal junctions are colored in lime green; thylakoid membranes in dark green; peptidoglycan layer and cytoplasmic membranes in gray; granules in yellow. Bar, 100 nm.

**Video S2.** Cryo-tomogram of a septal region of Δ*cse* mutant followed by segmentation, related to (Fig. 4b) and revealed a strong reduction of septal junctions. Septal junctions are colored in lime green; thylakoid membranes in dark green; peptidoglycan layer and cytoplasmic membranes in gray; granules in yellow. Bar, 100 nm.

**Video S3.** Cryo-tomogram of a septal region of Δ*cse* mutant followed by segmentation, related to (Fig. S4), which did not reveal any septal junctions. Thylakoid membranes are colored in dark green; peptidoglycan layer and cytoplasmic membranes in gray; granules in yellow. Bar, 100 nm.

